# Network topology in brain tumor patients with and without structural epilepsy- a prospective MEG-study

**DOI:** 10.1101/2022.12.01.518725

**Authors:** Barbara Ladisich, Stefan Rampp, Eugen Trinka, Nathan Weisz, Christoph Schwartz, Theo Kraus, Camillo Sherif, Franz Marhold, Gianpaolo Demarchi

## Abstract

It has been proposed that functional connectivity (FC) and network topology (NT) are altered in patients with glial brain tumors. So far there is no consensus on the pattern of these changes, furthermore data on FC in patients with brain metastases (BMs) as well as on the presence and absence of tumor-related epilepsy is lacking.

We aimed to analyze preoperative NT of newly diagnosed, singular, supratentorial glial brain tumors (GBTs) and BMs with and without structural epilepsy.

FC and NT derived from resting state magnetoencephalography (MEG) were compared between patients (PAs) and matched healthy controls (HCs), between GBTs and BMs as well as between patients with and without structural epilepsy. We investigated all patients, who met our inclusion criteria from February 2019 to March 2021. Clinical data was collected from patients’ electronic medical charts. We analyzed whole brain (wb) connectivity in six frequency bands, calculated three different network topological parameters (node degree (ND), shortest path length (L), clustering coefficient (CC)) and performed a stratification, where differences in the power were to be found. For data analysis we used Fieldtrip, Brain Connectivity Matlab toolboxes and in-house built scripts.

We consecutively included 41 patients (21 men), mean age 60.1ys (range 23-82), who were operated on at our institution. Tumor histology included high-grade gliomas (n=18), low-grade gliomas (n=4), dysembryoplastic neuroepithelial tumor (DNET, n=1), BMs (n=14) and others (n=4). Statistical analysis revealed a significant decrease for wb ND in patients compared to healthy controls in every frequency range analyzed at the descriptive and corrected level (p_1-30Hz_=0.002, p_γ_=0.002, p_β_=0.002, p_α_=0.002, p_θ_=0.024, p_δ_=0.002). Furthermore, at the descriptive level, we found a significant augmentation for wb CC (p_1-30Hz_=0.031, p_δ_=0.013) in PAs compared to HCs, which did not persist the false discovery rate (FDR) correction. There were no differences in the networks of glial brain tumors and metastases identified. However, we found a significant increase in wb CC in patients with structural epilepsy (p_θ_= 0.048), and significantly lower wb ND (p_α_= 0.039) at the uncorrected level.

Our data suggests that network topology is altered in brain tumor patients, which is in line with previous studies. Tumor histology per se might not influence the brain’s functional network, however, tumor-related epilepsy seems to do so. Longitudinal studies and in-depth analysis of possible factors and confounders might be helpful to further substantiate these findings.

## Introduction

Several symptoms observed in patients with brain tumors, such as attention deficits or reduced psychomotor speed, affecting quality of life, cannot be explained solely by the location of the lesion but might be explained by changes in the underlying neuronal network (Bartolomei et al., 2006).

The neuronal network is often characterized by means of functional connectivity and network topology (Bullmore and Sporns, 2009; Stam et al., 2009; C.J. Stam, 2010; Stam and van Straaten, 2012). Functional connectivity describes how information is transmitted locally as well as between different areas and is defined by statistical interdependencies of signals resulting from brain activity (Guggisberg et al., 2008; Stam et al., 2007). These signals can be measured by functional neuroimaging techniques such as functional magnetic resonance imaging (fMRI), or neurophysiological procedures, such as electroencephalography (EEG) or magnetoencephalography (MEG).

Network topology describes how the elements defining the network are arranged and how they interact. It is based on a mathematical graph theory that applies to various networks. This model has been increasingly applied in the fields of cognitive and clinical neuroscience, showing that functional and neuronal networks have so called “small-world” properties (Douw et al., 2011; Stam et al., 2009). Small-world networks are characterized by short path lengths (L) and high clustering coefficient (CC) (Barahona and Pecora, 2002). Douw et al. calculated a small-world coefficient (ratio between CC and L) from resting-state MEG and performed neuropsychological testing in healthy controls and found that higher small-worldness significantly correlated with better cognitive performance (Douw et al., 2011). A short path length indicates that nodes of a network relate to each other via short paths and high clustering describes that the probability of several nodes to form connections is high (Boccaletti et al., 2014). The node degree is another network topological parameter, which is an absolute value that specifies the number of connections of one single node (Vogel et al., 2021).

Among adult intraparenchymal brain tumors, brain metastases are the most common, followed by gliomas (Nayak et al., 2012). Even though prior studies already investigated the functional network of patients with glial brain tumors (GBTs), non-glial lesions and meningiomas, brain metastases (BMs) – to the best of our knowledge-have not yet been investigated in regard to their impact on the brain’s network (Bartolomei et al., 2006; Bosma et al., 2009; Derks et al., 2019, 2021; Douw et al., 2013; Semmel et al., 2021; van Dellen et al., 2012; van Nieuwenhuizen et al., 2018; Wang et al., 2010; Yang et al., 2022).

Even though low-grade gliomas (LGGs) and high-grade gliomas (HGGs) both derive from neuroepithelial tissue, they are different in terms of malignancy/ -clinical outcome, growth patterns, molecular signature, and protein expression (Louis et al., 2016). In contrast, BMs are secondary neoplasms that derive from completely different cell lines. Tumors from the lung, prostate, breast, melanoma, renal cell carcinoma most commonly spread to the brain (Nayak et al., 2012). The growth pattern of brain metastases is distinct from glial brain tumors, which can be visualized radiologically and intraoperatively. High-grade, as well as low-grade gliomas mostly infiltrate the brain parenchyma, whereas BMs usually only displace the surrounding tissue.

The authors of previous studies grasped to identify reasons for network changes in brain tumor patients. They proposed that histology and tumor grade may explain and directly impact the extent of network changes (Derks et al., 2019; van Dellen et al., 2012). It has been previously described that LGGs have a different influence on the network characteristics than HGGs, namely decreased synchronizability and reduced global integration (van Dellen et al., 2012). Further it has been found that patients with non-glial lesions, LGGs and HGGs show altered functional connectivity and impaired network topology, especially in the theta band, compared to healthy controls (Douw et al., 2010; van Dellen et al., 2012). These changes were also associated with impaired cognition and weren’t found to be explicable by the tumors’ anatomical location.

Another mechanism for network changes in brain tumor patients might be structural (symptomatic) epilepsy. Tumors causing structural epilepsy might have an even greater negative impact on the functional network compared to those that do not. It has been suggested that LGGs have a greater impact on network topology than HGGs (Bosma et al., 2009; Douw et al., 2013, 2010; van Dellen et al., 2012). This corresponds to the fact that LGGs also cause structural epilepsy more often than HGGs. Douw et al. further investigated tumor related epilepsy and found it to be interdependent with network characteristics and even to be related to protein expression (Douw et al., 2013). Previous MEG-studies on epilepsy found increased functional connectivity for focal and generalized epilepsy, as well as evidence for an increase in seizure frequency by augmented dissociation of the functional network (Douw et al., 2015; Li Hegner et al., 2018). So far, however there has not yet been any investigation published that compared the network topology of patients with versus without tumor related epilepsy.

Network changes, i.e., could be the cause or consequence of diseases. It might be lesions, like brain tumors, or pathological processes, like in Alzheimer’s/ Parkinson’s Disease or Epilepsy, that provoke alterations in functional networks (Douw et al., 2015; Engels et al., 2017; Hyder et al., 2021). Conversely, the distortion of the network could also generate the symptoms, or rather the diseases themselves.

Compared to the different impact of low-grade gliomas and high-grade gliomas on functional connectivity, we hypothesized that brain metastases would interact even more differently with their environment, respectively within the brain’s functional network, in comparison to the group of glial brain tumors.

And since previous studies found the FC in epilepsy to be clearly impaired, (Douw et al., 2015; Li Hegner et al., 2018; Rampp et al., 2021; Vogel et al., 2021) we now aimed to further investigate the effect of tumor-related epilepsy on NT.

## Study objective

The main objective of this study was to investigate whether network topology is altered in patients with brain tumors and whether histology or the presence of structural epilepsy have roles in these alterations.

We also aimed to confirm results from previous studies by applying current and most common network parameters and intended to add further information to the understanding of (pathological) network topology by investigating brain metastases.

We hypothesized that there is a relevant difference in network topology of patients with brain tumors compared to healthy controls, that BMs have a different influence on the brain’s network compared to GBTs and that structural epilepsy relevantly influenced network topology compared to patients without symptomatic seizures.

## Methods

### Study population

We included all consecutive adult patients who were referred to our department for neurosurgical resection or biopsy of a single, supratentorial tumor with MRI-radiological suspicion of either being a glial brain tumor or brain metastasis from 4^th^ of February 2019 to 21^st^ of March 2021.

We excluded patients who had undergone prior neurosurgery or radiotherapy of the brain, patients with unrelated psychiatric, neurological, endocrine comorbidities or implanted (non-titanium) metallic devices.

We matched every patient with a healthy control (HC) in gender and age (+/-5 years). The control group consisted of staff of the University Hospital Salzburg, as well as of students from the Paris-Lodron-University, Salzburg, Austria.

Both groups underwent a single 20-minute resting-state MEG recording. For co-registration with MEG data, a template MRI was used for HCs, whereas patients’ T1-pre- and post-gadolinium or T2- sequences were employed for the tumor group.

Clinical data was retrieved from patients’ electronic medical charts. In particular, the presence of structural epilepsy was recorded. In most patients with structural epilepsy, tumors had been initially detected due to the occurrence of seizures. Whenever this was unclear, an EEG was performed, and a neurological consultation was conducted.

The patient cohort was divided into two subgroups: glial brain tumors (GBTs) and brain metastases (BMs), and patients with structural epilepsy (PSEs) and patients without structural epilepsy (PNSEs).

### Ethics and dissemination

The study was reviewed and approved by the local ethics committee prior to recruitment (415-E/2412/6-2018). All eligible subjects gave their written informed consent to participate in this study. The Declaration of Helsinki was respected when conducting this study.

### MEG recordings

We recorded the brains’ magnetic signal at 1000 Hz (hardware filters: 0.1–330 Hz) in a standard passive magnetically shielded room (AK3b, Vacuumschmelze, Germany) using a whole head MEG (Neuromag Triux, MEGIN Oy, Finland). Signals are captured by 102 magnetometers and 204 orthogonally placed planar gradiometers at 102 different positions. All subjects wore nonmetallic clothes and had 5 head position indicator (HPI) coils attached on the glabella, forehead, as well as preauricularly bilaterally. They all underwent head shape acquisition prior to the recording, and the head shape points were later used for coregistering the anatomical MRI scan. During the measurement, subjects were seated and instructed to keep their eyes open. We periodically checked if subjects stayed awake via a camera in the magnetically shielded room. Every measurement consisted of one block of 20 minutes.

### Data analysis

To preprocess the data, we employed a signal space separation (SSS) algorithm that is implemented in the Maxfilter program (version 2.2.15), provided by the MEG manufacturer (Taulu and Simola, 2006). Here external noise from the MEG signal is removed (mainly 16.6 Hz train mains supply, and 50 Hz plus harmonics) and data is realigned to a common standard head position (*-trans default* Maxfilter parameter). This is performed across different blocks based on the measured head position at the beginning of each block using the five coils that have been placed before the measurement.

We used Brainstorm (Version 181008 (08-Oct-2018)) to further preprocess the data (Tadel et al., 2011). At first, we applied a high-pass filter at 0.1 Hz (6th order zero-phase Butterworth filter) to the continuous data. Subsequently, for signal space projection (SSP), continuous data were chunked in 10 s blocks (Uusitalo and Ilmoniemi, 1997). The resulting data was manually scrutinized to identify remaining eye blinks, eye movements, heartbeat and the 16 and 2/3 Hz train power supply artifacts and to create appropriate projectors. Finally, the artifactual components were projected out and the continuous data was segmented in 2s chunks. Then, data was again visually scrutinized to find bad epochs that were not corrected by the SSS and SSP combination, and further removed. An average of 15.4 (range 0-152.6) trials, corresponding to approximately 0.5 minutes were therefore removed, leaving roughly 19 minutes of artifact free data per subject (range 15-20 min) that was used for further analysis.

The subsequent steps were then performed using the *fieldtrip* toolbox (git version 20180124) (Oostenveld et al., 2011). At first, the data was downsampled to 300 Hz, to speed up the following computations of the source space connectivity and graph theoretical values.

Then the coordinate frame of the MEG data was coregistered to a template MRI scan in the case of healthy controls, and to T1-contrast enhanced or T2 anatomical images (depending on best visualization of the tumor) in patients, using the right and left preauricular points, nasion and the head shape points measured during preparation. A single shell source model was generated based on the segmented anatomical scan and a per subject source model with resolution 10mm was created (Nolte, 2003). Next, the data was Fourier transformed for each of the frequency bands of interest (‘broadband’=1-30 Hz, gamma=25-40 Hz, beta=15-25, alpha=8-15 Hz and theta=4-7, delta=1-4) was computed and used for a partial canonical coherence beamformer, with a noise covariance regularization factor of 5% (Oostenveld et al., 2011).

From the resulting source activity, for each position of the source model the component of the dipole moment projected onto the dominant direction was taken and subsequently used to compute the imaginary part of the coherence as an all-to-all whole brain voxel-wise connectivity metric. The 95th percentile of coherence values was used as a threshold for the construction of the adjacency matrix for computing node degree (ND), clustering coefficient (CC), shortest path length (L). All source, connectivity and network analyses were carried on using the fieldtrip toolbox (git version 20180124) (Oostenveld et al., 2011).

To minimize the spurious connectivity effects driven purely by the power difference, we decided to stratify the data to be sent to source space for connectivity and network analysis. For selecting the strata, two big pools containing the power spectral densities values were computed overall 2s blocks on the magnetometers, for the patients and for the healthy controls respectively. Then, the fieldtrip function *ft_stratify* (with the parameters method = ‘histogram’, equalbinavg = ‘yes’, numbin = 20) was employed to select the trials and equalize the two power spectral distributions, per each frequency band of interest.

### Surgery and histopathology

All decisions on neurosurgical treatment (consisting of either microsurgical tumor resection or biopsy) were agreed upon in the interdisciplinary neuro-oncological tumor board. Patients gave their written, informed consent prior to surgery.

Histopathological examinations (frozen section, formalin-fixed and paraffin embedded tissue, immunohistochemical and molecular genetic analysis) were performed according to the 2016 World Health Organization Classification of Tumours of the central nervous system, the current edition available during the inclusion period/ histological sampling (Louis et al., 2016).

### Statistical analysis

For statistical analysis we used Jamovi, Version 1 and R studio, Version 2022.02.1 (Şahin and Aybek, 2019; Wickham et al., 2019). As a dependent variable the maximum value of each network measure, per subject/patient and per frequency band of interest, was taken, that is node degree (ND), clustering coefficient (CC) and shortest path (L). We tested for differences in age in the patient population by means of Student’s t-test and Welch’s test. We designed a contingency table for one subgroup (PSE vs PNSE) -for which it was applicable- and calculated Fisher’s exact test. For the comparison between the groups 1) PAs vs HCs, 2) GBTs vs BMs and 3) PSEs vs PNSEs Shapiro Wilk test for normality was calculated. When the normality assumption was met, we performed independent Student’s T-tests, when it was violated, Mann-Whitney U tests were employed. Consequently, effect sizes were calculated by Cohen’s d or Rank biserial correlation (RBC), depending on the test employed. As customary, for p-values lower than 0.05 the null hypothesis was rejected, and the test was considered significant. We additionally provide the Bayes Factors (BF) to inform on the relative levels of evidence for the null/alternative hypotheses. At last, we corrected for multiple comparisons by calculating the false discovery rate (FDR).

Methods prior to statistical analysis are roughly summarized by a graphical depiction in Figure 1.

**Figure 1:**
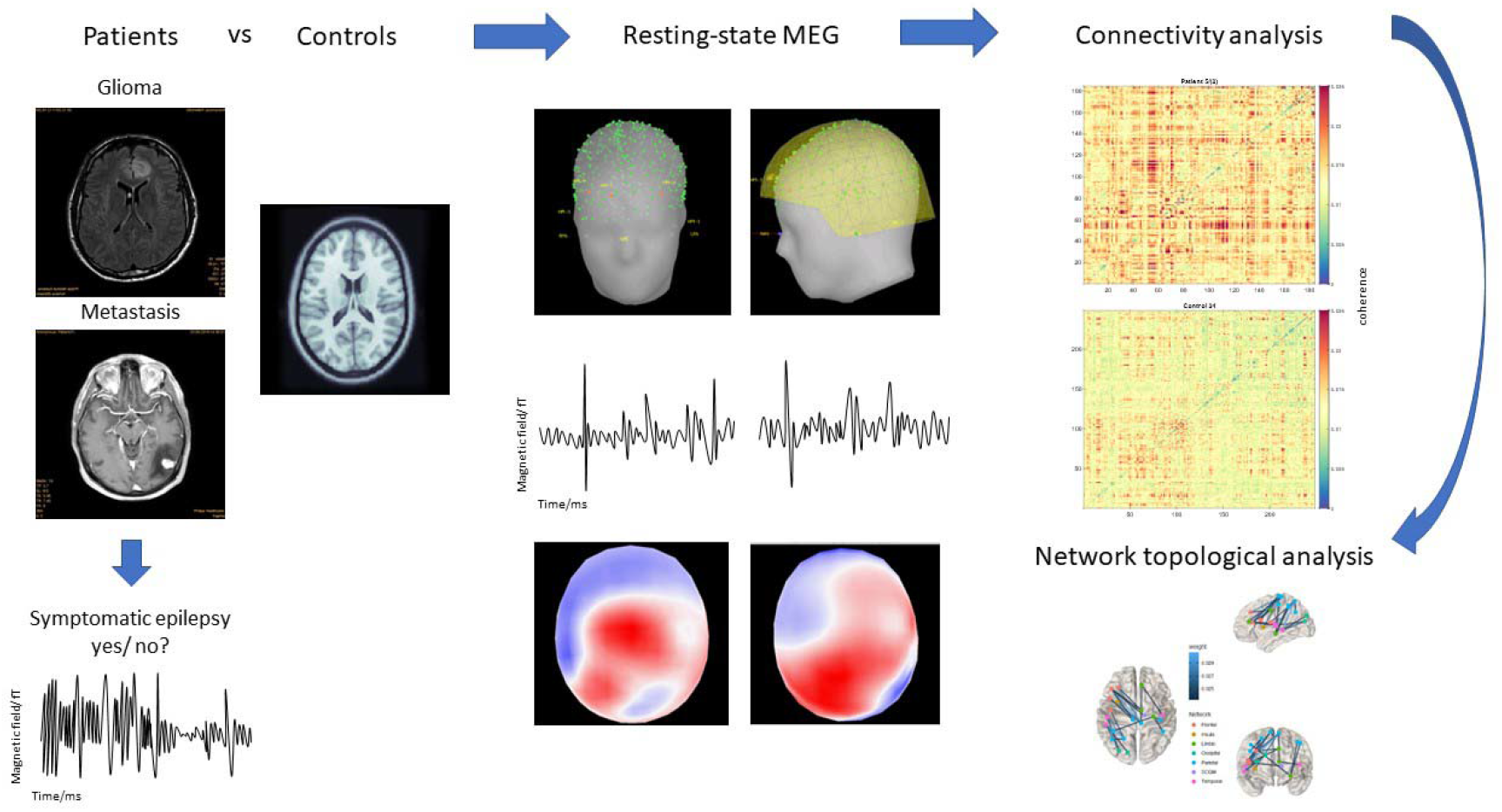
Summary of data collection and analysis. Patients with glial brain tumors and brain metastases were consecutively included and matched with healthy controls. Presence of structural epilepsy was recorded. MEG examination was performed several days preoperatively, whole brain connectivity was analyzed, resultant analysis of network topological parameters (node degree, shortest path length, clustering coefficient) was performed with Fieldtrip, Brain Connectivity Matlab toolboxes and in-house built scripts.

## Results

### Clinical data

A total of 42 patients were initially enrolled in the study. One patient diagnosed with acute disseminated encephalomyelitis by stereotactic biopsy had to be excluded from further analysis. Thus, the final study population consisted of 41 patients (21 men), as well as 41 healthy controls, who were matched in gender and age. Tumor histology included glial brain tumors (n=22), dysembryoplastic neuroepithelial tumor (DNET) (n=1), brain metastases (n=14), meningiomas (n=2) and primary CNS lymphomas (PCNSL) (n=2). The patient with DNET was investigated together with patients who had glial brain tumors, so that the final group of GBTs included 23 subjects.

None of the patients had undergone prior tumor treatment, besides corticosteroids or antiepileptogenic drugs at the time of MEG measurement.

Mean age in patients was 60.5years (+/- 15.2ys). T-Test and Welch’s Test revealed that there was no significant difference in age between patients with GBTs and brain metastases (p=0.260 and p=0.195 respectively).

A total of 27 patients were suffering from structural epilepsy: 15 with GBTs, ten with BMs and two with primary CNS lymphomas. Fisher’s exact test revealed no significant difference in the analysis of the contingency table.

Patient characteristics are shown in Table 1, 2 and 3.

**Table 1:** BM= brain metastases, HC=healthy controls, GBT= glial brain tumors, N= number, PCNSL= primary CNS lymphoma.

| **Characteristic** | **GBT** | **BM** | **Meningioma** | **PCNSL** | **HC** |
| --- | --- | --- | --- | --- | --- |
| N | 23 | 14 | 2 | 2 | 41 |
| Mean age (years) | 58.2 | 64.1 | 46 | 75.8 | 60.8 |
| Gender |  |  |  |  |  |
| Male | 14 | 6 | 0 | 1 | 21 |
| Female | 9 | 8 | 2 | 1 | 20 |

**Table 2:** DNET= dysembryoplastic neuroepithelial tumor (this patient was analyzed together with the GBTs), HGG= high-grade glioma, LGG= low-grade glioma.

| **Lesion type** |  | **N** | **Mean age** | **Male** | **Female** |
| --- | --- | --- | --- | --- | --- |
| **Glial brain tumors** | HGG | 18 | 60 | 11 | 7 |
|  | LGG | 4 | 46 | 3 | 1 |
|  | DNET | 1 | 21 | 0 | 1 |
| **Brain metastases** | Lung | 6 | 68 | 3 | 3 |
|  | Other | 3 | 59 | 1 | 2 |
|  | Breast | 2 | 53 | 0 | 2 |
|  | Renal | 2 | 68 | 1 | 1 |
|  | Melanoma | 1 | 70 | 1 | 0 |

**Table 3:** Presence of structural epilepsy in the patient cohort. Fisher’s exact test revealed no significant difference for the contingency table (p= 0.287). GBT= glial brain tumors; BMs= brain metastases; PCNSL= primary CNS lymphoma.

|  | **N with structural epilepsy** | **N without structural epilepsy** |
| --- | --- | --- |
| **GBTs** | 15 | 8 |
| **BMs** | 10 | 4 |
| **Meningiomas** | 2 | 0 |
| **PCNSL** | 0 | 2 |

### Network characteristics

We computed the resting state power spectral density (PSD) analysis for patients (PAs) and healthy controls (HCs) that revealed significant differences (p<0.001, RBC= 0.65, BF_10_=907). Therefore, to minimize the spurious connectivity effects driven purely by the power difference, we decided to stratify the data to be sent to source space for connectivity and network analysis. In the literature, there seems to be no standard way to summarize the whole brain network parameters, being mean, median, or maximum. Here, for simplicity, we report only the maximum values of the whole brain network parameters, but the usage of the mean and median delivers similar results (Hillebrand and Stam, 2019).

We found a highly significant lower node degree at the uncorrected level in patients compared to healthy controls (p_1-30Hz_ <0.001, d=-0.86, BF_10_=110.80; p_γ_=0.002, d=-0.72, BF_10_=19.35; p_β_<0.001, d=-1.13, BF_10_=6458.72; p_α_<0.001, d=-0.99, BF_10_=767.93; p_θ_=0.024, RBC=0.29, BF_10_=2.59; p_δ_<0.001, RBC=0.42, BF_10_=27.26) After the correction for multiple comparison, we still found a highly significant result in all six frequency bands as shown in Figure 1. Further analysis at the uncorrected level demonstrated significantly higher clustering coefficient (CC) (p_1-30Hz_=0.031, RBC=0.49, BF_10_=1.83; p_δ_=0.013, d= 0.32, BF_10_=9.07) in patients compared to healthy controls. However, the latter results of higher CC in patients showed a tendency but did not stand the correction (p_1-30Hz_=0.093, p_δ_=0.078).

For the comparison of the two tumor groups (glial brain tumors (GBTs) vs. brain metastases (BMs), we excluded the four patients with primary CNS lymphomas and meningiomas, respectively leaving 23 patients in the cohort of GBTs and 14 patients in the cohort of BMs. (Table 2). There was no discrepancy in power between the two groups (p=0.676, RBC= 0.09, BF_10_= 0.41) and in further network analysis we didn’t find a significant difference in any of the parameters and frequency bands.

We also compared patients with structural epilepsy against patients without structural epilepsy. For this analysis, we investigated all 41 patients-27 patients with versus 14 patients without structural epilepsy (Table 3). Here no power spectral difference between the groups was found (p=0.091, RBC=0.33, BF_10_= 0.66), so that no further stratification was needed. There was significantly higher maximum clustering (p_θ_= 0.048, RBC=0.38, BF_10_= 1.12), but significantly lower maximum degrees (p_α_= 0.039, RBC=0.40, BF_10_= 1.09) at the uncorrected level in PSE. However, after calculating the false discovery rate the two results did not stand the correction (clustering coefficient: p_θ_= 0.288; node degree: p_α_= 0.095).

Results are listed in Tables 4-7 as well as Figures 2 and 3.

**Table 4:** C= clustering coefficient; HCs= healthy controls; L= shortest path length; ND= node degree; PAs= patients. Maximum values are given. *Values that were found to be significant in later analysis.

| Frequency band | Network characteristic | Max |  |
| --- | --- | --- | --- |
|  |  | PA | HC |
| Broadband | ND | 227* | 287* |
|  | CC | 0.236* | 0.217* |
|  | L | 2.70 | 2.69 |
| Gamma | ND | 302* | 391* |
|  | CC | 0.270 | 0.265 |
|  | L | 2.78 | 2.75 |
| Beta | ND | 288* | 397* |
|  | CC | 0.281 | 0.269 |
|  | L | 2.83 | 2.82 |
| Alpha | ND | 300* | 398* |
|  | CC | 0.326 | 0.314 |
|  | L | 2.96 | 2.96 |
| Theta | ND | 348* | 399* |
|  | CC | 0.372 | 0.334 |
|  | L | 3.09 | 3.07 |
| Delta | ND | 367* | 454* |
|  | CC | 0.389* | 0.338* |
|  | L | 3.14 | 3.20 |

**Table 5:** BF10= Bayes factor_10_; C= clustering coefficient; d= Cohen’s d; HCs= healthy controls; L= shortest path length; ND= node degree; PAs= patients; RBC= Rank biserial correlation; *Values that were found to be significantly different between groups.

| Frequency band | Network characteristics | p-value | Effect size | BF<sub>10</sub> |
| --- | --- | --- | --- | --- |
| Broadband* | ND* | <0.001 | d=-.085 | 110.80 |
|  | CC* | 0.031 | d=0.49 | 1.83 |
|  | L | 0.623 | d=0.11 | 0.26 |
| Gamma* | ND* | 0.002 | d=-0.72 | 19.35 |
|  | CC | 0.875 | RBC=0.02 | 0.23 |
|  | L | 0.523 | d=0.14 | 0.28 |
| Beta* | ND* | <0.001 | d=-1.12 | 6458.72 |
|  | CC | 0.479 | d=0.15 | 0.29 |
|  | L | 0.711 | d=0.08 | 0.24 |
| Alpha* | ND* | <0.001 | d=-0.99 | 767.93 |
|  | CC | 0.705 | RBC=0.05 | 0.30 |
|  | L | 0.796 | RBC=0.03 | 0.23 |
| Theta* | ND* | 0.024 | RBC=0.29 | 2.59 |
|  | CC | 0.08 | d=0.39 | 0.89 |
|  | L | 0.717 | d=0.08 | 0.24 |
| Delta* | ND* | <0.001 | RBC=0.42 | 27.26 |
|  | CC* | 0.013 | RBC=0.32 | 9.07 |
|  | L | 0.218 | d=-0.27 | 0.45 |

**Table 6:** C= clustering coefficient; L= shortest path length; ND= node degree; PSEs= patients with structural epilepsy; PNSEs= patients without structural epilepsy. Maximum values are given. *Values that were found to be significant in later analysis.

| Frequency band | Network characteristic | Max |  |
| --- | --- | --- | --- |
|  |  | PSE | PNSE |
| Broadband | ND | 210 | 250 |
|  | CC | 0.214 | 0.215 |
|  | L | 2.70 | 2.68 |
| Gamma | ND | 299 | 351 |
|  | CC | 0.260 | 0.273 |
|  | L | 2.87 | 2.80 |
| Beta | ND | 291 | 343 |
|  | CC | 0.282 | 0.289 |
|  | L | 2.90 | 2.91 |
| Alpha* | ND* | 306* | 354* |
|  | CC | 0.325 | 0.340 |
|  | L | 2.99 | 2.97 |
| Theta* | ND | 370 | 373 |
|  | CC* | 0.400* | 0.361* |
|  | L | 3.19 | 3.15 |
| Delta | ND | 342 | 390 |
|  | CC | 0.357 | 0.363 |
|  | L | 3.16 | 3.11 |

**Table 7:** C= clustering coefficient; L= shortest path length; ND= node degree; PSEs= patients with structural epilepsy; PNSEs= patients without structural epilepsy; BF= Bayes Factor_10_; RBC= Rank biserial correlation (effect size); *Values that were found to be significantly different between groups.

| Frequency band | Network characteristic | p-value | Effect size | BF<sub>10</sub> |
| --- | --- | --- | --- | --- |
| Broad band | ND | 0.055 | RBC=0.37 | 2.22 |
|  | CC | 0.935 | d=-0.02 | 0.32 |
|  | L | 0.488 | RBC=0.14 | 0.37 |
| Gamma | ND | 0.079 | d=0.60 | 1.12 |
|  | CC | 0.581 | d=-0.18 | 0.36 |
|  | L | 0.148 | d=0.49 | 0.74 |
| Beta | ND | 0.078 | d=-0.60 | 1.12 |
|  | CC | 0.881 | RBC=0.03 | 0.33 |
|  | L | 0.923 | d=-0.03 | 0.32 |
| Alpha* | ND* | 0.039* | RBC=0.40 | 1.09 |
|  | CC | 0.470 | d=-0.24 | 0.39 |
|  | L | 0.712 | d=0.12 | 0.34 |
| Theta* | ND | 0.956 | RBC=0.01 | 0.32 |
|  | CC* | 0.048* | RBC=0.38 | 1.12 |
|  | L | 0.494 | d=0.23 | 0.38 |
| Delta | ND | 0.067 | RBC=0.35 | 0.66 |
|  | CC | 0.786 | d=-0.09 | 0.33 |
|  | L | 0.694 | RBC=0.08 | 0.41 |

**Figure 2:**
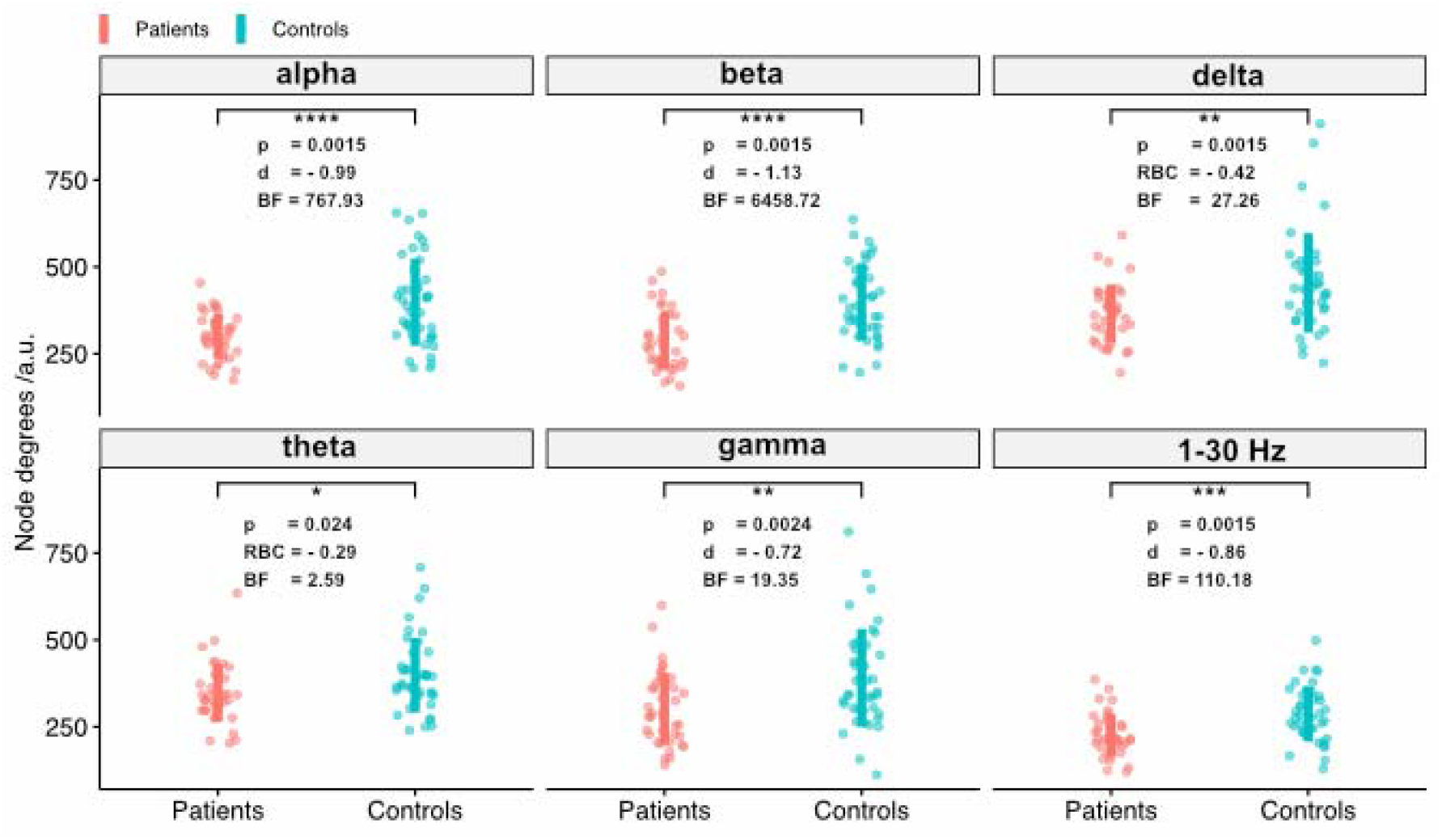
T-test or Mann-Whitney U test comparison (depending on normality distribution) between patients (red) and healthy controls (blue) revealed significantly lower node degree in patients at the corrected level (p1-30Hz =0.0015, pγ=0.0024, pβ=0.0015, pα=0.0015, p_θ_=0.024, p_δ_=0.0015).

**Figure 3:**
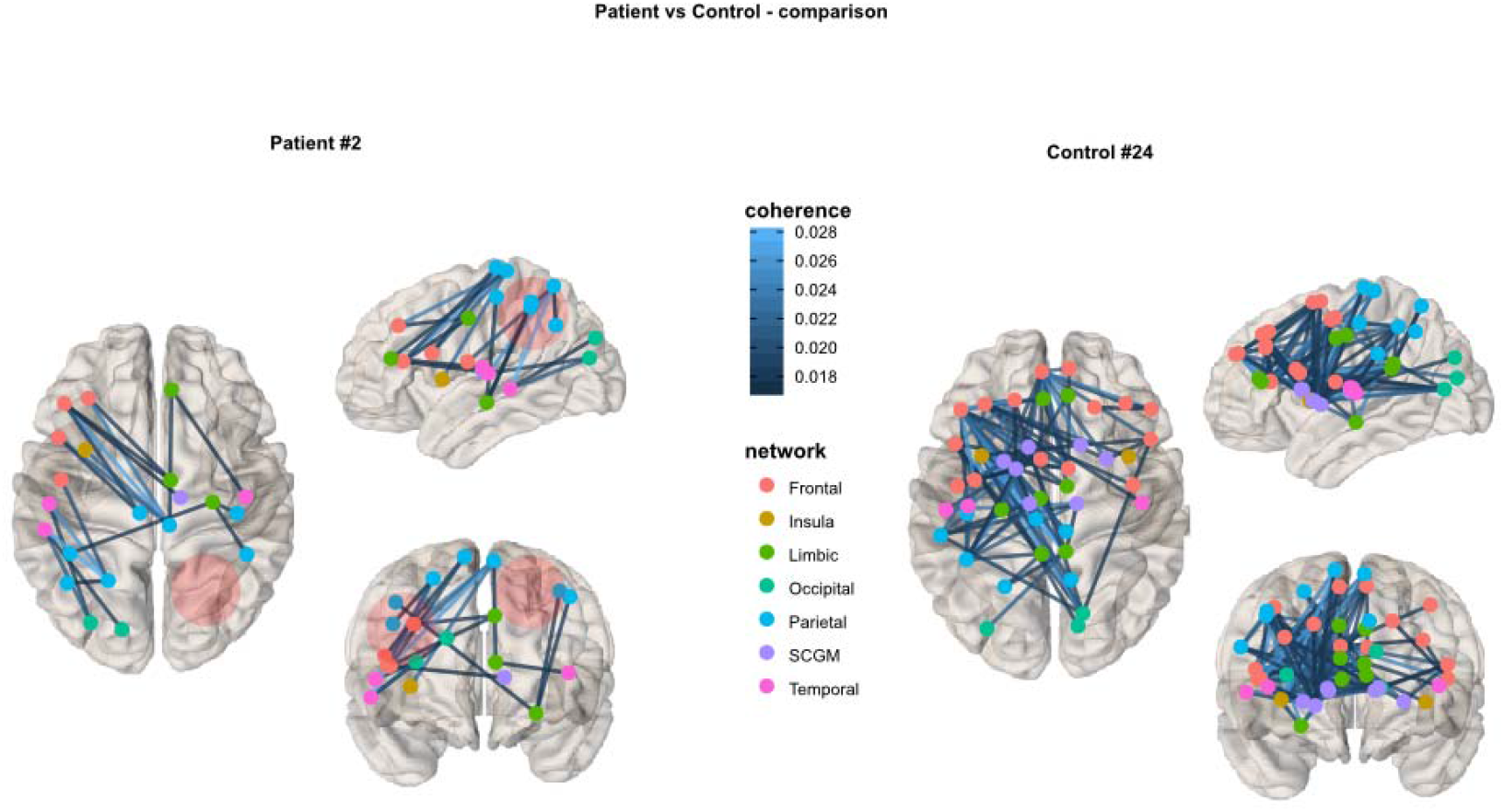
Representative coherence plot thresholded at 75% of the maximum respective coherence of a patient and its matched control: on the left is patient 2, who was diagnosed of a metastasis in the right parietal lobe, on the right is its matched control depicted.

## Discussion

It was previously described that different kinds of neurological disorders, such as brain tumors lead to deviations in the brain’s functional network (Bartolomei et al., 2006; Cho et al., 2021; De Baene et al., 2017; Derks et al., 2021, 2019; D’Souza et al., 2021; Douw et al., 2013; Huang et al., 2014; Hyder et al., 2021; Na et al., 2018; van Dellen et al., 2012; Wang et al., 2010).

In this prospective study we investigated 41 patients with brain tumors and 41 matched healthy controls. We were able to identify significant differences in the network topology of patients compared to healthy controls, as well as between patients with and without structural epilepsy. Unexpectedly, we found no differences in the functional networks of patients with glial brain tumors compared to brain metastases.

### Comparison of patients and healthy controls

In the comparison of patients and healthy controls we found a significantly lower node degree in patients in all frequencies analyzed. This result is also consistent with the definition of node degree since the impact of the tumor on the functional network will impair the formation of connections and therefore lead to lower degree networks (Vogel et al., 2021).

We found an increased clustering coefficient in brain tumor patients at the descriptive level, but this result did not stand up to correction for multiple testing.

The clustering coefficient is a value that is defined by the probability that neighbors of a hub are connected to another. It is an essential characteristic of the small-world concept that describes an optimally functioning network (Bullmore and Sporns, 2009; Douw et al., 2011; Stam et al., 2009). This result appeared unusual at first sight, since an intact network is assumed to have high clustering and we would have expected these parameters to be decreased in a pathologically disturbed network, as it is the case in a patient with a brain tumor. Several previously conducted studies have also described increased clustering in brain tumor patients (Bosma et al., 2009; van Dellen et al., 2012; Derks et al., 2021). Bosma et al. found a higher clustering coefficient within the theta band in patients with low-grade gliomas compared to healthy controls (HCs) (Bosma et al., 2009). They contextualized this result in the setting of other neurological diseases such as major depression, autism disorders or Alzheimer’s disease, for which previous investigations have found similar network changes. Also, Derks et al. described higher global clustering in glioma patients compared to HCs. They further investigated this finding by differentiating various factors (e.g., age, histology, local and global clustering) but despite this, could not find any relation to increased clustering of any of these aspects (Derks et al., 2021). As such, it was concluded that the probable cause of the increased clustering lay with diffuse, whole brain alterations in these patients. The augmentation might also be an expression of reactive changes of the network to the tumor. This assumption seems at least conclusive from a clinical point of view since lesions also provoke morphological and immunochemical reactions in the surrounding parenchyma that can be displayed radiologically and by the laboratory.

### Comparison of patients with glial brain tumors and brain metastases

We further performed two analyses solely within the patient cohort. The first one between the patients with glial brain tumors (GBTs) versus brain metastases (BMs). For this we excluded four patients, due to their histological diagnoses-two showed PCNSL, the other two had meningiomas. The latter two patients were primarily included since both had an oncological history and from a neuroradiological point of view it had not been possible to rule out a BM. There was one patient, in which we had the initial suspicion of a LGG, which then turned out to be a DNET. Although DNETs are a distinct tumor entity, and do not belong to the group of gliomas, we kept this case included within our cohort of GBTs, because of clinically similar behavior.

Since data on the impact of BMs on the brain’s functional network is lacking, we chose to investigate those patients in this context. We expected metastases to show different patterns of network changes, due to the completely different type of tumorigenesis and growth patterns. However, this hypothesis cannot be supported by our data since no difference in the functional networks of GBTs and BMs was found, - neither in the variables nor the frequencies analyzed.

Previous studies had found different network patterns in HGGs and LGGs, which had led to the conclusion that histology plays an important role in network changes (Derks et al., 2019; van Dellen et al., 2012). Derks et al. even found significant differences in the immunohistochemical profile of tumors with lower alpha functional connectivity (FC) in IDH-wildtype gliomas compared to IDH-mutated gliomas. Nonetheless they did not find differences in FC in the theta band between the two groups, and there was also no difference in the alpha band in the comparison of classic histological grading between grade II/III and IV (Derks et al., 2019).

Our results might be limited by the fact that the two tumor groups were not exactly equally distributed. This is due to the prospective setting of this study and the fact that single (supratentorial) brain metastases are rare.

Still, it is a noticeable finding, and if metastases and gliomas are indeed comparable regarding their impact on network topology, then the main factors may be the mass effect or growth rate, rather than histology. Metastases usually show less invasive growth patterns compared to intrinsic brain tumors but may cause substantially more mass effect due to common extensive peritumoral edema. Therefore, if these tumors do not lead to different changes in the functional network, histology might not be an explanation for the disturbed network.

### Comparison of patients with and without structural epilepsy

Another possible mechanism of the distortion of the functional network could be the presence of seizures. We therefore also conducted a comparison between patients with and without structural epilepsy. Here, we found significantly higher clustering (CC) and lower node degree (ND) in PSEs at the uncorrected level. After calculating the FDR these results did not retain significance.

However, we still want to discuss the finding of the lower ND in patients with structural epilepsy, since it stands in contrast to previous studies that presented data from patients with drug resistant focal epilepsies (Rampp et al., 2021; Vogel et al., 2021). In their data functional connectivity and node degree was augmented in patients with epilepsy compared to healthy controls. One explanation for this contradiction might be that tumor-related structural epilepsy has a different impact on the network than non-tumor-related focal epilepsy. Patients with epilepsy mostly suffer from their disease for many years, in contrast to patients with brain tumors who are primarily structural with seizures and shortly after receive diagnosis and treatment for the lesion. In the situation of long-lasting epilepsy, with many events of seizures, the network might distort and adapt completely differently than in structural epilepsy.

In this study, we applied graph theoretical measures since they have proven to be a valuable tool to characterize the network topology of the brain (Bullmore and Sporns, 2009; Heimans and Reijneveld, 2012; Reijneveld et al., 2007; Stam and Reijneveld, 2007). Some of the most current network topology parameters are clustering coefficient, path length and node degree (Boccaletti et al., 2014; Bosma et al., 2009; van Dellen et al., 2012; Vogel et al., 2021; Wang et al., 2010). Until now there is only one review on the topic by Semmel et al., who found that the methodology is often not comparable and that the results are sometimes even contradictory between different publications (Semmel et al., 2021).

Therefore, we also carried out functional connectivity and network topology calculations in source space, compared to other previous studies that computed the connectivity measures in sensor space only. It is crucial to measure in source space, especially in patients with morphological brain pathologies (Stufflebeam, 2011).

Whenever we found significant results in a network topological parameter, the increase/ decrease compared to healthy controls was concordant in the other frequency bands. To us, this supports the reliability of the results.

On the contrary, another study found, for example, an increase of synchronization and clustering in theta, but a decrease in beta (Bosma et al., 2009). They argued that there was more small world organization in lower frequency bands than in higher ones.

It seems that this aspect needs further investigation.

### Strengths and limitations

This is a prospective, experimental, single-center study that aims to further increase the neurophysiological understanding of how brain tumors disturb the brain’s functional network. The study population might be small but is comparable to other studies (Bartolomei et al., 2006; Bosma et al., 2009; Derks et al., 2021, 2019; van Dellen et al., 2012; van Nieuwenhuizen et al., 2018; Wang et al., 2010). Due to the prospective setting, and because we included patients in a consecutive fashion, subjects were not exactly equally distributed in the two groups (glial brain tumors vs brain metastases and patients with vs without structural epilepsy). On the one hand, this hampers the comparableness between groups, but on the other side we can provide higher data quality and therefore higher significance.

We put a focus on rigorous methodology for best possible comparability with previous and future studies. Therefore, we only investigated patients with a newly diagnosed, singular, supratentorial mass lesion, suspected to be a primary brain tumor or brain metastases. Further we arranged MEG measurement for all patients a few days, and up to a maximum of two weeks prior to surgery. In this way, we aimed to prevent possible influences of previous treatment, the presence of new lesions or additional tumor growth on the results of network analysis. A defined timing of MEG with respect to treatment can help to define standards in network analysis.

To ensure the significance of our results, we corrected for multiple comparisons, as we compared 6 different frequency bands for every network parameter. After this, the comparison of node degree in patients and healthy controls was still highly significant. Admittedly the increased clustering coefficient in patients was not found to be persistently significant, but there was still a noticeable tendency to be seen. The corrected results of the comparison between patients with and without structural epilepsy were not found to be significant. Nevertheless, we mentioned these results, because they differ from previous publications and we expect this distinction to be an interesting aspect for future investigations. So far, there exist only seven other studies on the topic of functional connectivity and network topology measured by MEG in brain tumors (Bartolomei et al., 2006; Bosma et al., 2009; Derks et al., 2021, 2019; van Dellen et al., 2012; van Nieuwenhuizen et al., 2018; Wang et al., 2010). Although previous studies tried to draw conclusions from the influence of histology on network alterations, BMs have not been investigated, though, from a pathohistological point of view, they differ the most from primary brain tumors. We aimed to close this gap through our analysis, however our results did not support the role of histology in network disruption.

## Conclusion

We provide data of a prospective study on patients with brain tumors with and without structural epilepsy compared to healthy controls. We found significant differences in several network parameters in the broad band, as well as in all five different frequency bands. We did not find differences in the functional network of primary brain tumors compared to brain metastases, but there were significant differences at the descriptive level between patients suffering from structural epilepsy compared to those that did not.

Originating from the results of this study the extent of network disruption in patients with brain tumors seems to be influenced more by the presence of structural epilepsy, than by the histology.

We advocate further prospective studies on even larger cohorts of patients. Especially the differentiation of patients with structural epilepsy, patients suffering from epilepsy and the comparison to healthy controls could be essential.

## Abbreviations and acronyms

BF: Bayes factor
BM: brain metastasis
CC: clustering coefficient
DNET: dsyembryoplastic neuroepthelial tumor
EEG: Electroencephalography
FC: Functional connectivity
FDR: False discovery rate
fMRI: Functional magnetic resonance imaging
GBT: Glial brain tumor
HC: Healthy control
HGG: High-grade glioma
HPI: Head position indicator
IDH: Isocitrate dehydrogenase
L: Shortest path length
LGG: Low-grade glioma
MEG: Magnetoencephalography
MRI: Magnetic resonance imaging
ND: Node degree
NT: network topology
PA: Patient
PCNSL: Primary CNS lymphoma
PSD: Power spectral density
PNSE: Patients without symptomatic epilepsy
PSE: Patients with symptomatic epilepsy
RBC: Rank biserial coefficient
SSS: Signal space separation
SSP: Signal space projection
wb: Whole brain

## Appendices

## Acknowledgements

To all my colleagues, friends and family, and the students from Lodron University who took their time to participate in this study as healthy controls.

To Ciara O’Sullivan for providing language help.

To Österreichische Krebshilfe Salzburg for their research funding, that partly supported this project. The financial support was used for data collection and analysis. There was no involvement in writing of the report, or in the decision to submit the article for publication

## Data and code availability

The MATLAB scripts used to perform the analysis are available at one author’s (GD) repository (github.com/gdemarchi). Down sampled (to 100 Hz) version of the healthy controls raw data will be available at OSF.io upon final publication. The sharing of the patients’ data must be instead discussed upon request on a per-case basis with the CDK/PMU Salzburg Ethics committee.

